# A study of a diauxic growth experiment using an expanded dynamic flux balance framework

**DOI:** 10.1101/2022.05.16.492165

**Authors:** Emil Karlsen, Marianne Gylseth, Christian Schulz, Eivind Almaas

**Author notes:** These authors contributed equally to this work (Marianne Gylseth, Christian Schulz).

## Abstract

Flux balance analysis (FBA) remains one of the most used methods for modeling the entirety of cellular metabolism, and a range of applications and extensions based on the FBA framework have been generated. Dynamic flux balance analysis (dFBA), the expansion of FBA into the time domain, still has issues regarding accessibility limiting its widespread adoption and application, such as a lack of a consistently rigid formalism and tools that can be applied without expert knowledge. Recent work has combined dFBA with enzyme-constrained flux balance analysis (decFBA), which has been shown to greatly improve accuracy in the comparison of computational simulations and experimental data, but such approaches generally do not take into account the fact that altering the enzyme composition of a cell is not an instantaneous process.

Here, we have developed a decFBA method that explicitly takes enzyme change constraints (ecc) into account, decFBAecc. The resulting software is a simple yet flexible framework for using genome-scale metabolic modeling for simulations in the time domain that has full interoperability with COBRA toolbox 3.0. To assess the quality of the computational predictions of decFBAecc, we conducted a diauxic growth fermentation experiment with *Escherichia coli* BW2523 in glucose minimal M9 medium. The comparison of experimental data with dFBA, decFBA and decFBAecc predictions demonstrates how systematic analyses within a fixed constraint-based framework can aid the study of model parameters.

Finally, in explaining experimentally observed phenotypes, our computational analysis also demonstrates the importance of non-linear dependence of exchange fluxes on medium metabolite concentrations and the non-instantaneous change in enzyme composition, effects of which have not previously been accounted for in constraint-based analysis.

## Introduction

Computer models are invaluable tools in capturing and systematizing new knowledge, especially for the complex phenomena found in biology. Genome-scale metabolic models (GEMs) are computational models that compile information about the entirety of known metabolic functions in a given organism or cell type. GEMs typically contain a listing of genes, enzymes, and reactions, and relationships of dependence between these. For biochemical reactions, information about substrate, product, and stoichiometry is included in their model representation, i.e. the consumption and production rates for the involved compounds. Based on how compounds participate in different reactions, it is possible to infer a metabolic network: a bipartite network connecting the reactions and metabolites [1]. An additional central component in GEMs is the representation of a biomass objective function (BOF), a pseudo-reaction which represents the metabolites needed for the cell to reproduce. Since it is a key component of these models, the BOF has recently been the target of increased interest in the field [2–6].

At their best, computational models help identify inconsistencies in our current understanding of the phenomena they model, predict novel system behavior or connections, and aid in the design of experiments [1, 7–10]. As the use of these models becomes more popular and necessary to improve systems-level understanding of metabolism, so should their ease of use and interpretation. Due to the formulation of the GEMs and the assumptions of steady-state, mass conservation, and optimality of an objective (commonly chosen to be the BOF), the calculation of system-wide flux-states (measured in millimoles per hour per gram of cell dry weight (mmol h^−1^ gCDW^−1^) [1]) can be performed using standard tools for constraint-based linear optimization [1]. This allows for very rapid arrival at an optimal solution, even for large networks containing thousands of reactions. The aforementioned analysis- and calculation-steps are called flux balance analysis (FBA) [1].

In the years since its inception, FBA and related approaches have given rise to a number of derivatives and modifications [11]. Two modifications that in particular improve the utility of FBA are (1) dynamic flux balance analysis (dFBA) and (2) enzyme-constrained flux balance analysis (ecFBA). In dFBA, the goal is to simulate the interaction between the organism’s metabolism and the environment over time [7, 12]. In the ecFBA approach, additional constraints are applied to the flux distribution to account for the fact that the proportion of active enzymes in a cell is only a fraction of the cell mass, and these enzymes have a finite capacity to catalyze biochemical reactions [13, 14].

While first introduced in 1994 [7], dFBA was first explicitly formalized in 2002 [12]. This formalization emphasized two main approaches: the dynamic optimization approach (DOA), and the static optimization approach (SOA) [12]. The essential difference between the two methods is that in DOA, the simulation is solved for a single interval of time (often the total duration of interest) which determines the optimal strategy, whereas in SOA, a regular FBA problem is solved for a sequence of time intervals, and the effects from one interval are propagated to the next until the final interval is solved [12]. Effectively, this means that for the same environment over the same total timeframe, DOA is likely to produce a sequence of metabolic operations that results in a higher sum score for whatever objective it is optimized for than SOA, as it is able to “plan ahead” e.g. by rationing some to-be-limiting nutrients. One might therefore speculate that, in general, DOA is likely a better predictor for an organism well adapted to a predictable environment, while SOA is likely a better predictor for an opportunistic organism in a novel or unpredictable environment. The method for solving the problems also differ, as the SOA can be solved using simpler algorithms, being a series of regular (linear) FBA problems [12]. No matter the kind of dFBA formulation, it can be a useful tool for simulating the conditions in a system such as a bioreactor over time [8]. dFBA, however, faces a few key challenges, such as unrealistically rapid shifts in flux distributions, and the occasional numerical issue [15].

Several software packages have been built that enable the performance of simple dFBA [15–21], but many are either difficult to acquire [15, 19], have become unavailable [17], or only contain parts of a solution [16, 20], thus requiring decisions to be made on the part of the modeler that may produce different results from the same underlying data. Such decisions are always a part of modeling, but in the case of dFBA, they are not usually explicit, leading to a number of different solutions to the same basic problems.

A notable exception is the modelling framework COMETS, which hosts a range of functionalities and provides interfaces for several popular programming languages to its open-source code, although most of that code is implemented in Java [18, 21]. Overall, the tools for performing dFBA are generally aimed at expert users who are either able to improvise the required coding themselves or have very specific aims.

Most biochemical reactions need to be catalyzed by enzymes to proceed at a physiologically relevant rate (or at all). As an enzyme both has a finite rate of catalysis and a mass, this places a constraint on the possible productivity of a given amount of cells. While there are a few different approaches to incorporate this as a limitation in FBA [9, 13, 22], they all spring from the above assertions and assumption that this carries additional, meaningful constraints on the flux distribution in a metabolic network. Here, we collectively refer to these approaches as enzyme-constrained FBA (ecFBA). Besides the sound arguments underlying the ecFBA approaches, they also improve the ability of the constraint-based framework to reproduce some phenomena observed experimentally that are outside the capability of basic FBA without the addition of extensive and somewhat arbitrary constraints. An example is the modelling of overflow metabolism, where glucose is incompletely metabolized, thus yielding a much lower amount of energy per glucose molecule, yet a higher amount of energy per time [9, 23]. While the ecFBA approaches are useful in many situations, these models are data-hungry and require extensive information on enzymes associated with different reactions, namely their mass and the turnover numbers [9]. As these enzyme-constrained models can be somewhat cumbersome to build, requiring extensive retrieval processes from databases such as SABIO-RK [24] and BRENDA [25], efforts have been made to facilitate the process through automation and more standardized protocols [9, 14, 26].

A common application of FBA is to study the interaction between a given organism and its environment over time, such as in the case of a batch fermentation [11]. While it is not strictly necessary to run a dFBA simulation of the entire process to do so, batch fermentation provides a natural testing ground for assessing the accuracy of both the GEM and the model of the fermentor itself, as well as testing the validity of the different assumptions at work. It is also a display of a GEMs predictive powers that might be easier to grasp than more domain-specific analysis methods such as phenotype-phase-plane analysis [27]. Due to the potential profits of improving the efficiency of industrial batch production of added-value bio-molecules, and the requirements for model fidelity in order to do so, a natural step then becomes to combine the simpler-to-interpret aspects of dFBA with the increased fidelity of ecFBA [28–31], a combination we will call decFBA in the following. The DOA approach requires added theoretical as it increases the complexity of both formulating and solving the mathematical problems involved. In the case of SOA, it can be implemented in a straightforward way. One of the key merits of ecFBA is that it improves a GEM’s ability to reproduce certain metabolic phenomena crucial to accurate description of cellular behavior, such as overflow metabolism [9].

Also required for accurate simulation is the BOF, which is defined as a pseudo-reaction, listing the amount of different (pseudo-) metabolites needed per hour to produce a growth rate of 1h^−1^. As a model of this type operates in a gCDW^−1^ capacity, this becomes the transient amount of different metabolites needed for 1 gCDW to produce 1 gCDWh^−1^, without explicitly considering the compounding effects of exponential growth [1]. As the BOF serves a central role in GEMs, determining the overall flux distribution by being the objective the entire system it is trying to optimize for, it should receive special care and attention. While the traditional approach has been to approximate it based on data compiled from literature, often from different strains or conditions, the subject has been the target of increasing interest in recent years [2–6]. Yet, much work remains to be done, both in terms of establishing protocols and analysis techniques [6], and in terms of a theoretical framework for integrating the resulting data [5] before we can reasonably expect BOFs to allow accurate predictions in varying and novel environments. Due to its central role in metabolism, the BOF is of special importance when trying to quantitatively predict metabolic behavior. Thus, the simulation of a concrete situation can be a useful test for verifying a given GEM biomass composition. This is especially true when looking at parameters such as terminal biomass, where the amount of biomass produced in a given fermentation should be predictable in a simulation using a GEM containing an accurate BOF.

In the following, we expand decFBA using the SOA approach by explicitly adding constraints on how rapidly the mass fraction occupied by enzymes can be reallocated, named enzyme change constraints (ecc). We demonstrate how this has significant effects on modeling dynamics by showing its impact on GEM behavior during simulation of generic diauxic growth on glucose. Finally, we proceed to test the impact of the ecc on a GEM’s ability to recreate a specific fermentation experiment, and how the ecc can help explain observed behavior.

## Materials and methods

### Batch fermentation

A carbon-limited batch fermentation of *E. coli* BW25113 (hereafter *E. coli*), was carried out in a 3 L Eppendorf NewBrunswik BioFlo 115 bioreactor in batch setup, using minimal glucose media as specified in S1 File. Glycerol stock solution samples prepared from a single colony were grown in 250 mL baffled shake flasks overnight at 37 °C and 200 rpm shaking in glucose minimal M9 medium. The fermentors containing 1.5 L of sterile medium were then inoculated with 40 mL of culture. The fermentation parameters were 37 °C, pH 7, and 40 % dissolved oxygen (DO). During setup, the pH electrodes were calibrated with pre-mixed solutions of pH 4 and 7. The pH was prevented from dropping below 7 through automated addition of NaOH (4 M). The DO electrodes were calibrated to 0% by flushing the electrode for 20 min with nitrogen gas, and to 100% by running the reactor with the medium and filtered air inflow of ≈600mL min^−1^ at 500 rpm stirring until equilibrium. Stirring was then coupled to DO, ensuring an oxygen saturation in the medium of ≥ 40%, with 200 rpm and 1000 rpm as lower and upper bounds. Analysis of the off-gas composition and flow-rate was performed using a ThermoScientific Prima BT Benchtop MS, which was calibrated before each run.

Sampling was performed every 30 minutes for optical density (OD) measurements and medium analysis, and every 30-60 minutes for dry weight. The OD was measured using a spectrophotometer at a wavelength of 600 nm by sampling 5 mL. An aliquote of the sample was diluted with purified water such that the measured OD was between 0.2 and 0.6. The medium samples were taken as 2 mL aliquotes from the OD samples. They were spun at 6700 RCF for 2 min, the supernatant was sterile filtered (0.2 μm polyethylensulfon syringe filters) and stored at −20 °C until it was analysed. For dry weight measurements 10 mL were sampled, washed (centrifuged and redissolved) once with 5 mL 0.9% NaCl before being centrifuged again and redissolved in purified ion-free water, and added to a pre-dried and -weighted aluminum pan. The pans were completely dried (for ≥36 h at 120 °C) before being weighed again.

### Medium analysis

The quantification of medium constituent concentrations was performed by NMR, using ERETIC2 [32]in the Bruker TopSpin 4.0.8 software. The protocol is based on Søgaard *et al.* [33] and was further developed.

The samples were thawed and mixed 1:10 D_2_O with 0.75% TSP (purchased from Sigma), before 600 μL of the resulting solution was added to a 5 mm 7 inch NMR tube, in which it was analyzed using a 600 MHz (14.1 T) Bruker NMR spectrometer running with the H_2_O + 10% D_2_O solvent setting and water suppression (^1^H NMR, noesygppr1d). The acquisition parameters were: 4 dummy scans, 32 scans, SW 20.8287 ppm, O1 2820.61 Hz, TD 65536, TE 298.0 K, D1 4 s, AQ 2.621 439 9 s. P1 was calibrated for each sample to ensure accurate quantification. A creatine solution (70 mM in D2O-TSP solution) was used as an external standard, quantified using the singlet at ~ 3 ppm. Compounds responsible for peaks in the spectra were identified based on reference^1^H-NMR spectra available in the Human Metabolome Database (HMDB) [34–36], as well as the software Chenomx [37] and published literature by Fan [38]. Note that, glucose quantification was not based on the α-Glucose doublet at ~ 5.2 ppm, as these peaks can be affected by the water suppression sequence [39]. Instead, we used the quartet at ~ 3.2 ppm for this quantification (reference HMDB compound ID HMDB0000122).

### GEM, COBRA and code availability

The GEM used for all modeling was the *E. coli i*JO1366 [40] model, or more specifically, its enzyme-constrained modification *i*JO1366* as presented by Bekiaris [14]. The coding and modeling was performed with MATLAB 2020a, using version 3.0 of the COBRA Toolbox for MATLAB [20], and the linear program solver for version 2.7 of the GNU Linear Programming Kit (glpk) solver (https://www.gnu.org/software/glpk/). Using the glpk solver, the **feasTol** parameter had to be lowered from the default 1*e* − 6 to 1*e* − 9 to prevent numerical issues with the algorithm placing limits on enzyme reallocation.

The code produced and used in the paper is available in a supplementary zip file S3 File, and on GitHub, at https://github.com/emikar/FermentationSim. Anyone is permitted and encouraged to freely copy the code and implement their own modifications as needed. Details for the software can be found in supplementary file S2 File.

### From dFBA to decFBAecc

In order to test the effect of constraining enzyme allocation at different stages, we first ran a model equivalent to standard dFBA, then added total enzyme constraints (decFBA), and finally added enzyme change constraints (decFBAecc). We selected a simple test scenario: a 1 L reactor with an initial biomass of 0.1 gCDW growing on 10 mM glucose with bounds on glucose and oxygen uptake rates as specified in Table 1.

**Table 1.** Iterative change in constraints from dFBA to decFBAecc. The parameters that are subject to change are the total fraction of mass available for enzymes (ec; ggCDW^−1^) and the amount of enzyme in terms of total mass that can be replaced per hour (ecc; ggCDW^−1^h^−1^).

| Approach | ec | ecc |
| --- | --- | --- |
| dFBA | 5000 | - |
| decFBA | 0.0948 | - |
| decFBAecc | 0.0948 | 0.01 |

First, to get a regular dFBA equivalent, we used a version of *i*JO1366* with the protein-pool constraint increased to allow arbitrarily high fluxes, effectively rendering the enzyme constraint inactive, and thus performing a standard dFBA. Second, we ran a version of the same GEM where the protein-pool constraint remained at its default setting, thus performing standard decFBA. Third, and finally, we ran a version of the model where the rate of enzyme redistribution was constrained to a plausible rate value, performing what we have called decFBAecc. For all of the simulations, we set the O_2_ maximum uptake rate to 15 mmol gCDW^−1^ h^−1^ and the glucose maximal uptake rate to 10.5 mmol gCDW^−1^h^−1^. The constraints are listed in Table 1, and the outcomes are given in the Results and discussion section.

### Reproducing fermentation results

The complete simulation setup with all required configuration and model files can be found in S3 File and on the project GitHub page, https://github.com/emikar/FermentationSim. The outcomes of using the software are examined in the Results and discussion section.

The well-known phenomenon of overflow metabolism [41] is relatively straight-forward to reproduce using basic FBA, but is better captured using a model which has had properly tuned enzyme constraints, such as the GECKO method [9]. As the AutoPACMEN [14] package is streamlined to allow the addition and tuning of enzymatic constraints to a GEM, its proof-of-concept, the *i*JO1366* *E. coli* model, was considered a suitable candidate for this work. Before tuning, all exchange reaction bounds were set to ±1000 mmol gCDW^−1^h^−1^.

In order to reproduce the results from the batch fermentation, the following tuning steps were required:

1. As micronutrients are not supposed to be limiting in the given situation, the various ions and trace minerals were set to not be exhausted from the medium.
2. As stirring was coupled to DO in the fermentation, oxygen was also set to not be exhausted from the medium.
3. The model-encoded uptake rates of glucose and oxygen were too high; these are closely linked, and as such the oxygen uptake was limited to a maximum of 15 mmol gCDW^−1^ h^−1^.
4. The following byproducts were excreted by the iJO1366* model in addition to, or instead of, acetate, and were therefore constrained to zero (both isomers where applicable), in the following order: lactate, dihydroxyacetone, ethanol, and pyruvate.
5. Finally, the excretion rate of acetate was adjusted until it best matched the observed concentrations, resulting in a maximal acetate excretion of 2 mM gCDW^−1^h^−1^ (see further discussion of this in the Results section).

The order in which these steps were carried out, along with the acetate excretion feedback relation discussed below and presented in a more systematic overview in Table 2.

**Table 2.** Model parameters for reproducing fermentation. The constraints that change through the iterative modifications in our attempt to recreate the experimental observations made during the fermentation. These correspond to the differences between the different panels in Fig. 3. The various values given for lb refer to the lower bounds on the corresponding exchange reaction in the COBRA model, while the ub refer to the upper bounds. The exchange reactions are: ox, oxygen; lac, lactate; dha, dihydroxyacetone; ac, acetate; pyr, pyruvate. Where applicable, both stereoisomers are affected by the changes. The ”ac feedback” parameter is described below.

| Iteration | ox lb | lac ub | dha ub | ac ub | ac lb | pyr ub | ac feedback |
| --- | --- | --- | --- | --- | --- | --- | --- |
| A | -1000 | 1000 | 1000 | 1000 | -1000 | 1000 | No |
| B | -15 | 0 | 0 | 1000 | -1000 | 1000 | No |
| C | -15 | 0 | 0 | 2.2 | -10 | 0 | No |
| D | -15 | 0 | 0 | 2.2 | -10 | 0 | Yes |

The medium composition was transcribed from the medium recipe (given in S1 File) into a CSV file where the concentrations of the different constituent compounds are given in mmol L^−1^. As implied by the tuning steps above, only the concentrations of glucose and ammonium were actually imposed as limits during the simulations, of which only glucose was exhausted, and therefore, had its concentration becoming relevant.

In order to get more clearly measurable signatures for the NMR medium analysis, higher concentrations than what is typical were used during the fermentation. An interesting side effect of this was that, the concentration of acetate seemed to reach saturation earlier than predicted by the computational simulations. This appears to be a known phenomenon, which is linked to thermodynamic feedback in the Pta-AckA pathway [42].

In order to capture this dynamic in a simple way, a formula for the upper bound on acetate excretion was devised based on figure 4a by Enjalbert et al. [42] and added to the model, as given in Eq. 1:

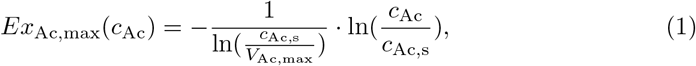

where *Ex*_Ac,max_ is used as the upper bound on acetate excretion, and *c*_Ac_ is the concentration of acetate in the environment. *V*_Ac,max_ = 2 is the max excretion rate for acetate, while *c*_Ac,s_ = 13 is the external acetate concentration at saturation. Fig. 1 shows a comparison of Eq. 1 with data from Fig. 4a, Enjalbert *et al.* [42].

**Fig 1.**
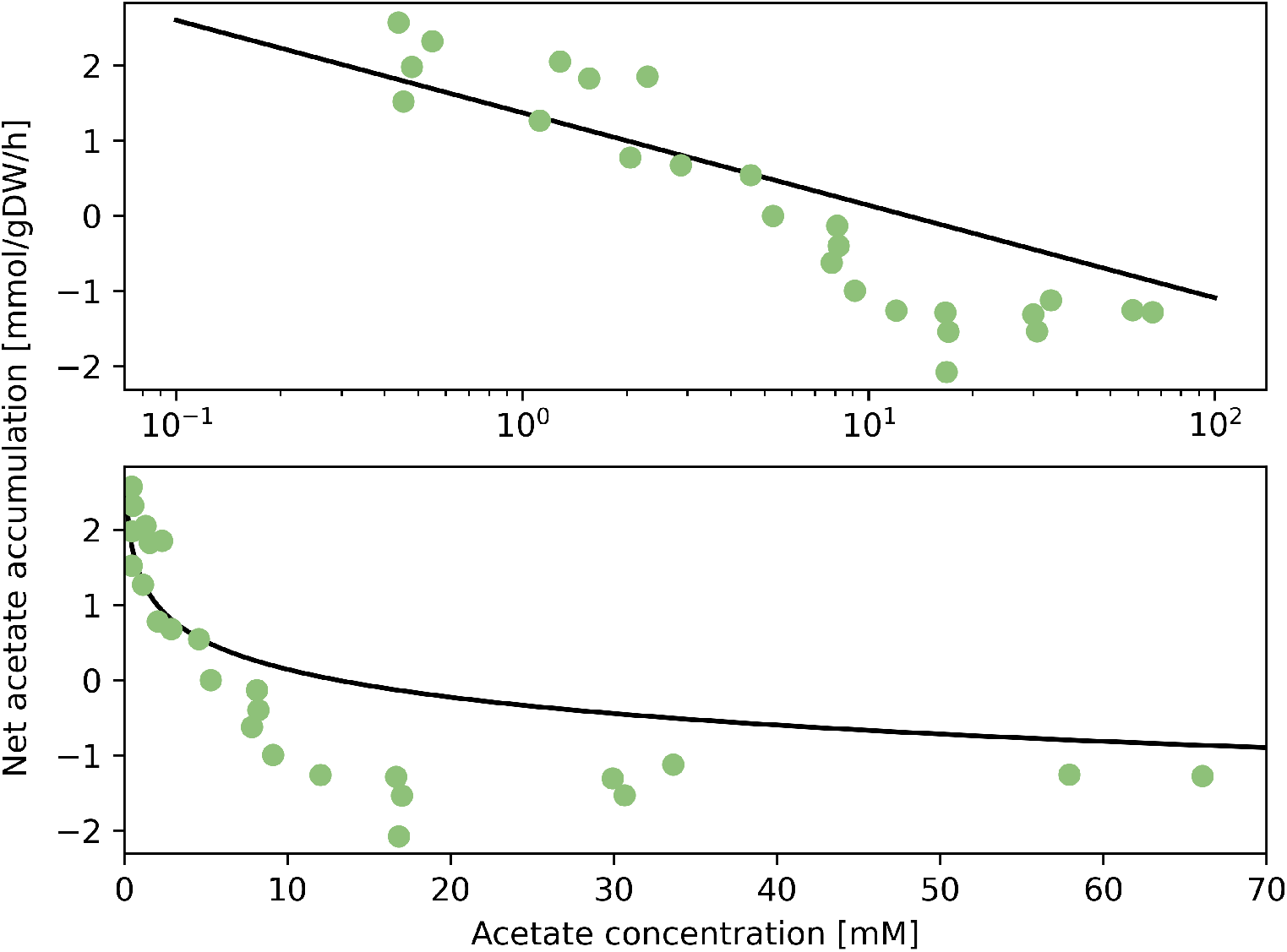
Acetate excretion feedback relationship. Assessment of Eq.1 (solid line) ability to match data (filled circles)from Enjalbert *et al.* [42] Fig. 4a. The fit is acceptable in the target range *c*_Ac_ < 10.

### Modeling a time-constrained change in enzyme composition

In order to infer enzyme compositions, the flux through each reaction was multiplied by the corresponding bottom element of the sMOMENT S-matrix and then multiplied by −1. This step corrects for the fact that the bottom row is all-negative to be balanced by the positive coefficient connecting them to the positive protein composition constraint. As the bottom element gives the molecular weight of the corresponding enzyme divided by the enzyme’s turnover number, multiplying by the flux through the reaction catalyzed by that enzyme yields the total amount of mass currently dedicated to that enzyme per gDW. This can be seen from Eq. 6 in [14]:

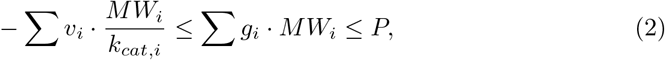

which implies:

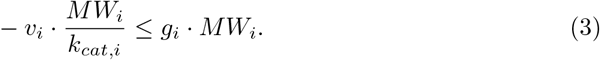

This leads to:

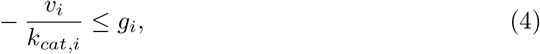

or, for our purposes, the following assumption:

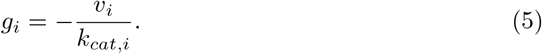

Using this equation, it is possible to keep track of the enzyme composition of the cell.

We apply an ecc rate limit by storing the enzyme composition as a separate vector variable, in terms of mass dedicated to the enzymes for each reaction. Subsequently, the ecFBA problem is solved once without enzyme composition change limits, then the implied enzyme composition is checked against the former composition. If the difference is smaller than or equal to the amount allowed for that time step (as calculated by multiplying the number of hours in the time step by the enzyme change limit given in gCDW h^−1^), the enzyme composition vector is updated to reflect the new composition. If, however, the difference is greater than the limit for that time step, the enzyme composition is calculated by weighted linear interpolated between the former state and the desired state, and a new ecFBA problem is solved with upper bounds on the flux values corresponding to the new enzyme composition.

Expressed mathematically, we have the enzyme composition vector of the current time step *m_t_*, giving the protein mass dedicated to each enzyme in gCDW:

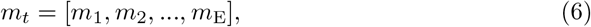

where E is the number of enzymes in the vector, which for the purposes of the sMOMENT model is inflated to the number of (uni-directional) fluxes in the model.

For each time step, the composition vector of the former time step *m*_*t*–1_ is compared to the optimal composition vector *m*_o_ acquired by solving a new ecFBA problem, thus giving the difference vector *m_d_*:

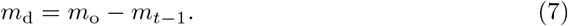

If the sum of the positive elements in this difference vector is smaller than the enzyme composition change constraint *γ* for the current time step, then:

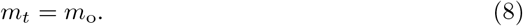

Otherwise, the value for *m_t_* will be calculated by weighted linear combination. First we find the proportion of the change, *α*, permitted by the composition change constraint *γ*:

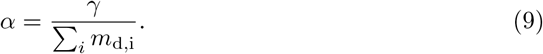

Next, we acquire the new *m_t_* by weighted linear combination of *m_o_* and *m*_*t*–1_:

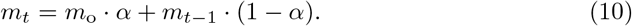

This option can be activated in the configuration file by setting the enzyme composition change limit for a given simulation. If set to −1, it allows the enzyme composition to change freely on every step without tracking, while set to any positive value (including 0), it constrains the change in enzyme composition to that number of gCDWh^−1^. Any other number will limit the rate of change in enzyme composition to that fraction of total biomass per gCDW per hour. E.g. setting it 0.1 will allow 0.1 grams of enzyme to be replaced per gCDW per hour, which for the standard enzyme allocation setting of the sMOMENT model *i*JO1366* of 0.0948 would amount to replacing 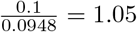, or 105%, of total enzyme per hour.

## Results and discussion

### Gradually constraining a dFBA model

In Fig. 2 we show the effects of adding different constraints to the dFBA formulation, thus transitioning from standard dFBA to a decFBA approach, to a situation with enzyme composition change constraints (decFBAecc).

**Fig 2.**
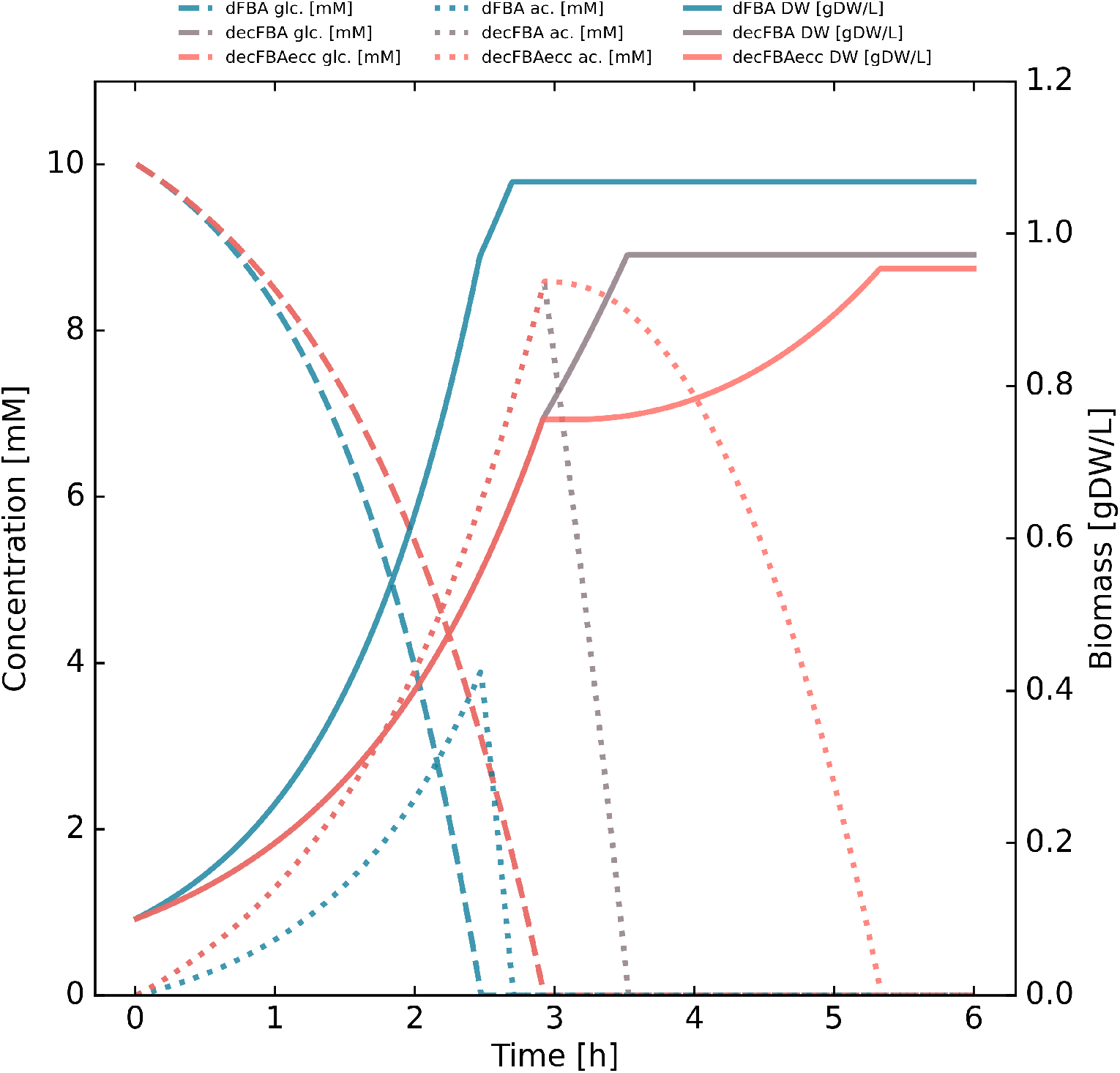
Transitioning from standard dFBA to decFBAecc. Most reactions are unconstrained, but glucose uptake is limited to 10.5 g gCDW^−1^ h^−1^, and oxygen uptake is limited to 15 g g CDW^−1^ h^−1^. In the dFBA case, the *i*JO1366* model has been adjusted with an arbitrarily large protein pool constraint (5000 g gCDW^−1^ of enzyme, which allows every flux to reach 1000 mmol gCDW^−1^ h^−1^) to produce standard dFBA results; glucose is rapidly depleted, while acetate is accumulated before being rapidly depleted as well. In the decFBA case, the *i*JO1366* model is simulated without modifying the protein pool constraint (0.0948 g gCDW^−1^ of enzyme), allowing slightly slower growth; glucose is depleted slower than in the dFBA case, and more acetate accumulates before being depleted at a similar rate. In the decFBAecc case, the protein pool constraint is the same as default setting, but the rate at which the enzyme composition is allowed to change has been limited to 1% per hour (as opposed to being instantaneous); the trajectories are identical until the metabolism shifts from growth on glucose to growth on acetate, at which point there is a short lag phase.

**Fig 3.**
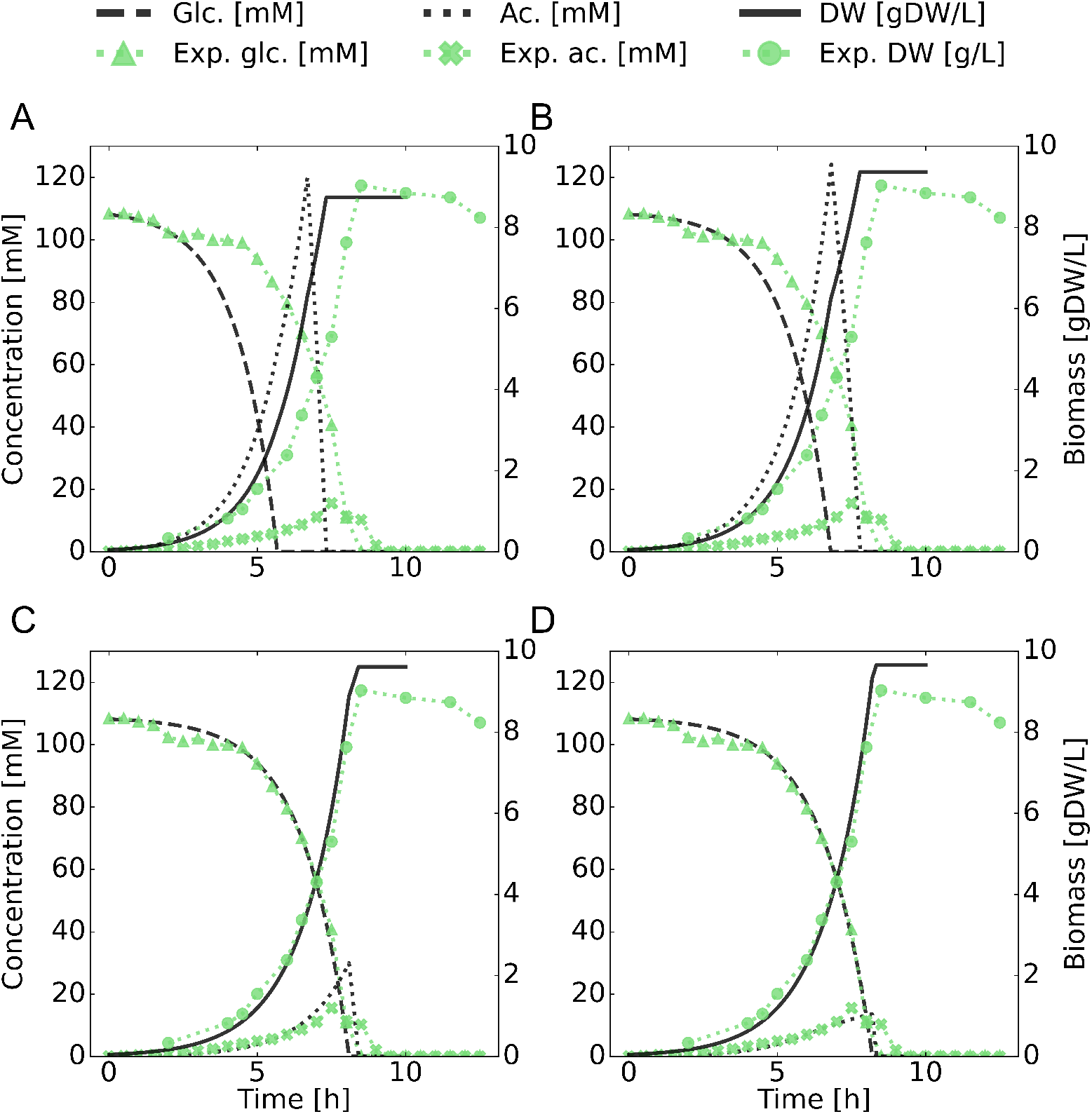
Model parameter adjustment. Comparison of modeling results with measurements throughout successive adjustments of base sMOMENT model parameters. As can be seen, several export reactions need to be blocked, and several rates limited in order to reproduce realistic behavior, even with the total enzyme pool constraints. The order and nature of adjustments are listed in Table 2. In short: panel A is *i*JO1366 out-of-the-box, panel B is oxygen uptake-limited with lactate and dihydroxyacetone export blocked. Panel C has acetate exchange bounded and pyruvate excretion blocked, while panel D has acetate excretion feedback enabled.

**Fig 4.**
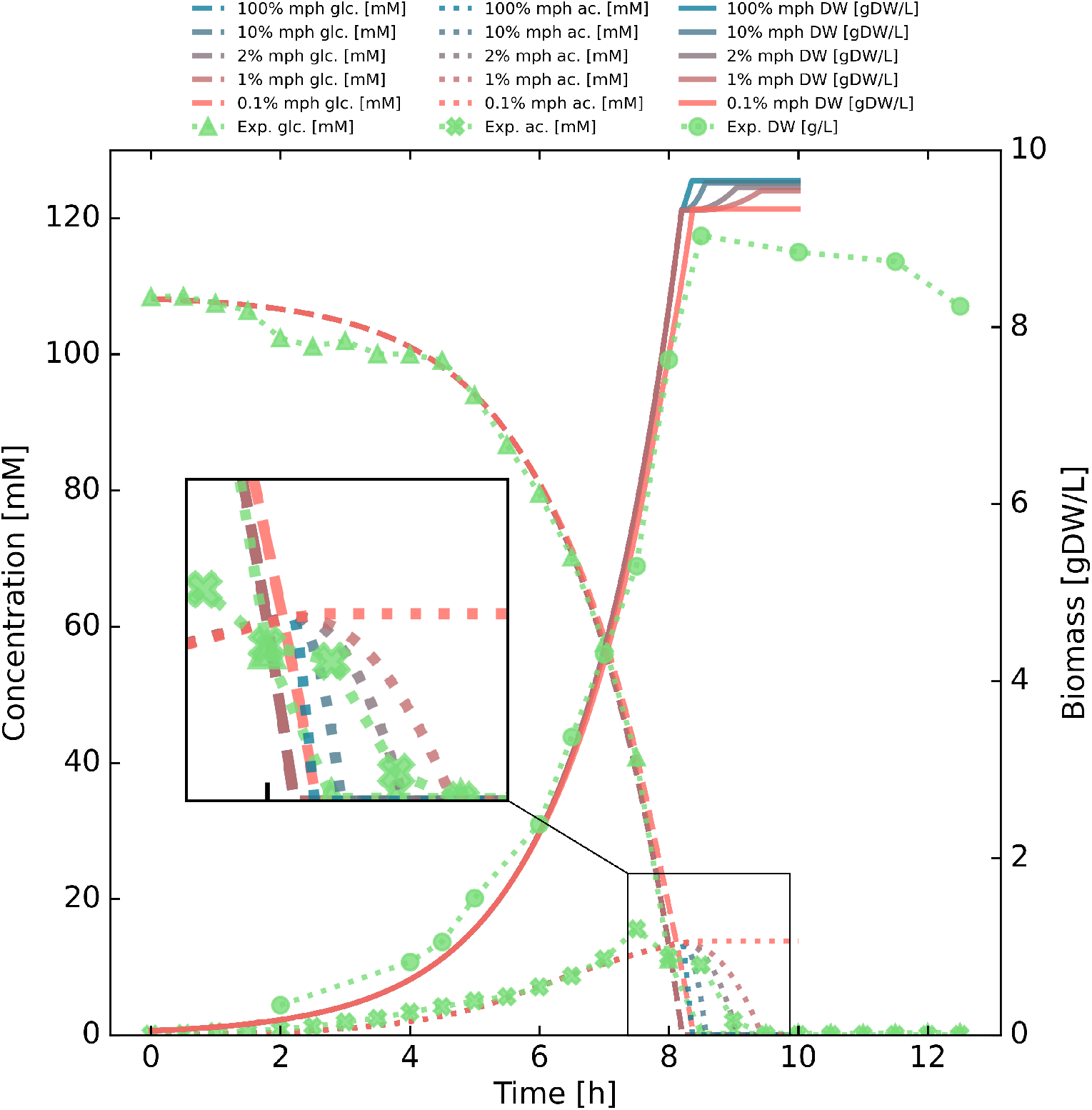
Enzyme composition change rate adjustment. Comparison of modeling results with experimental measurements when varying constraints on rate of change of enzyme composition. The rate is given as percentage of total cell mass (not enzyme mass) per hour (denoted mph in the legend) that can be reallocated. The rate of change of enzyme composition has a dramatic effect on the transient behavior of the model during diauxic growth, and a rate of 2% mph appears to best match the data measured from the fermentation experiment.

The application of these is described in detail in the Results and discussion section. The first parameter set (”dFBA”, Table 1) corresponds to standard dFBA, realized through increasing the sMOMENT protein pool constraint from 0.0948 to 5000 g gCDW^−1^ of enzyme. Here, glucose is rapidly depleted as the amount of biomass grows and acetate is excreted, before the model fully depletes the glucose and then also rapidly depletes the acetate.

The second parameter set (”decFBA”, Table 1) corresponds to decFBA as realized by running the sMOMENT approach unmodified in a dFBA setting. Here, glucose is consumed more slowly than for simple dFBA simulation, and acetate accumulates at a much more higher rate. This is typical when enzyme constraints are added, as incomplete metabolism of glucose through glycolysis produces more energy per time per amount of enzyme than does the full aerobic breakdown through TCA cycle and electron transport chain. Once glucose is depleted, the acetate concentration quickly follows suit at a similar rate to what is the result in the dFBA case.

The third parameter set (”decFBAecc”, Table 1) corresponds to decFBAecc as realized by running the sMOMENT model unmodified in a dFBA setting while constraining the amount of enzyme mass that can be reallocated per time. The resulting curves seem identical to the decFBA case until glucose is depleted. This is expected since the only difference is the rate at which enzyme can be reallocated, and both decFBA and decFBAecc are initialized at optimal enzyme composition for the consumption of glucose. Following this, the acetate depletion rate builds up gradually, as the biomass curve under decFBAecc displays a lag phase. The final biomass concentration is lowered due to the application of these constraints reducing the total conversion efficiency of glucose to biomass.

Overall, we find that constraining the rate at which enzyme mass can be reallocated has an important and profound impact on the dynamics of cell metabolism. While the impact in terms of terminal biomass is small between decFBA and decFBAecc, the impact of the timing at which the terminal biomass is reached is significant, which has implications for microbial production of compounds where a change in growth conditions often is implemented as a trigger for production of a compound in question. In conclusion, the inertia in the cells’ change in enzyme composition appears to warrant further investigation.

### Batch fermentation

During the exponential growth phases, growth rates were measured to 0.66 h^−1^, while the terminal biomass values were measured to 8.2 gCDW L^−1^. The experimental results, including sampling history and raw data from the fermentor’s monitoring software and off-gas analyzer are given in the S4 File.

The fermentation data are plotted in Figs. 3 and 4, where they are compared with different simulation results. As can be seen, the fermentation follows a typical timeline for diauxic growth of *E. coli* on glucose [7], with the depletion of glucose and concurrent build-up of acetate, which subsequently is depleted. The concentrations of the various organic compounds in the medium were determined using the NMR protocol outlined in the Materials and methods section. Of note is the observation that biomass per volume starts to drop off after peaking, likely due to starvation and cell death. Since the measurements of biomass are actual dry weight measurements, the drop-off is not a result of simple change in cell morphology.

### Adjusting model parameters

Fig. 3 shows the effects of the sequential adjustment of model parameters as we attempt to capture the specific metabolic behavior displayed in our experiment. The adjustments are listed in the Methods section and briefly in Table 2.

The base decFBA computational modeling, as seen in Fig. 3 panel A, uses the *i*JO1366 model. It depletes glucose much too fast and produces an acetate peak that is much too high. Terminal biomass is not too far off, however. In Fig. 3 panel B, oxygen uptake has been limited to a more realistic value (see Table 2), which caused excretion of lactate and dihydroxyacetate. As these were not observed experimentally, they have been blocked. Overall, this results in a slower glucose depletion and a higher acetate peak, without an appreciable change in terminal biomass. In Fig. 3 panel C, acetate excretion and uptake are limited to more realistic values (see Table 2) [42], which initially caused excretion of pyruvate, which was subsequently also blocked, as no appreciable amounts were detected in the medium. This last step has as a consequence that growth, glucose, and acetate curves align much better with experimentally measured values, indicating that the entire system’s behavior is sensitive to the maximal rates of acetate exchange. Finally, in Fig. 3 panel D, we show the results of adding an expression for negative feedback from external acetate concentration on acetate export (see the Materials and methods section for details) that is motivated by research by Enjalbert and coworkers [42]. The resulting concentration curve for external acetate appears to align even better with our experimentally measured values.

### Consequences of the enzyme change constraint

Next, we investigated the effect of applying a constraint on rate of change in the cells enzyme composition (Fig. 4), as formulated by the decFBAecc (see Materials and methods). The consequences of applying the constraint on the rate of enzyme reallocation are most apparent in the depletion of external acetate, and, to a lesser extent, in the terminal biomass. The more constrained the reallocation of enzymatic mass becomes, the slower acetate is depleted, simultaneously lowering the terminal biomass. This is to be expected, as the optimal enzyme distribution for utilization of glucose is not the same as the optimal enzyme distribution for utilization of acetate. In this particular case, a rate of replacing 2% of total biomass worth of enzymes per hour appears to produce an excellent fit with experimental data, though this rate is 1) likely to be highly dependent on possible rate-limiting components in the enzyme reallocation process, and 2) has a lower bound limited by how rapidly gene expression can occur.

## Conclusion

In this work, we have presented a computationally robust and flexible framework for simulating fermentation with dynamic flux balance analysis. We use this framework to make iterative changes in computational constraints applied to a simple example of diauxic growth on glucose. This approach illustrates the consequences of adding constraints on the rate of reallocation of enzyme. Based on these findings, we perform a simple fermentation experiment for *E. coli* as a case study, which we attempt to recreate using our computational simulation framework.

We have two primary findings. First, we find that applying constraints on the rate of enzyme mass reallocation, in addition to applying constraints on total mass allocation to enzymes, produces markedly different modeling results. The rationale for the new constraints are sound, and they do appear to allow a model to better capture experimental observations. Maybe the most easily observable consequence, is that they may markedly change the timing at which maximum biomass is reached during diauxic growth, which could have important implications for industrial production of added-value compounds. They do not, however, markedly change the terminal amount of biomass in the considered case, meaning their utility in verifying a BOF may be limited.

Second, we strengthen the notion that non-linear feedback dynamics on acetate excretion are necessary in order to capture the behavior of *E. coli* in environments with high initial concentrations of glucose. This agrees with the findings of Enjalbert et al. [42], and has important consequences for modeling of metabolism at high external concentrations of export products.

## Supporting information

**S1 File. The carbon-limited medium used in the fermentation**. An Excel file detailing the nutrient composition of the medium used in the fermentation.

**S2 File. Manual for the simulation program for Matlab**. A PDF manual containing specifications and user instructions for the fermentation simulation program written in Matlab.

**S3 File. Zip file of project source code**. A zip file containing the Matlab source code.

**S4 File. The fermentation data**. Excel file detailing the fermentation, including protocol and sampling history.

## Notes

https://github.com/emikar/FermentationSim

### Competing Interest Statement

The authors have declared no competing interest.

