## Supplemental File 2 for "A study of a diauxic growth experiment using an expanded dynamic flux balance framework"

### User manual accompanying: A study of a diauxic growth experiment using an expanded dynamic flux balance framework

Emil Karlsen

15.04.2022

This manual is a companion piece to the article: "A study of a diauxic growth experiment using an expanded dynamic flux balance framework". Here, the file structure hierarchy necessary for the program to run will be outlined and illustrated, along with the program structure and flow. Additionally, some examples are provided of how the software can be modified to represent other situations.

The simulation performed here is reminiscent of textbook performance of dFBA, but is configured in a clearly defined and structured fashion, where reconfiguration does not necessitate adding or altering lines of code. This design choice was made for several reasons: 1) it allows customization while ensuring all files and processes are loaded/called in the correct order, 2) it provides a formalized set of specifications that could be transferable between different software suites and programming languages, and 3) it may allow inexperienced programmers to modify the program settings more easily.

#### File structure

In order to run the program, certain files need to be in place, organized in the correct structure. Aside from the model files themselves, every file involved is in the CSV format. A hierarchical overview of the folders in the right hierarchy can be seen in Fig. 1. Any given simulation will only call on the data found in a single file in each subfolder, apart from the **input/media/** folder, from which several media may be called for a given fermentation. Note that a single call to `setup_simulation` may run several simulations, and may therefore call files from several subfolders, but will only call a single file from **input/configuration/** per received function call.

The folder contents are listed in alphabetical order:

- **input:** Folders containing the various inputs to the program. These are all looked up from the `setup_simulation` script.
    - **additions:** Different addition schemes. These list how much of the different media (which are retrieved from the **input/media/** folder) are added, and at what times. Only one such file is called for a given simulation.
    - **complex_rules:** Mathematical relationships between decFBA model variables and constraints. Contains a **constraint** column indicating the constraint to be modified, a **rule** column indicating the type of relationship, and a number of columns of parameters (**k1**, **k2**, ... , **kn**) for the relationship.
    - **configuration:** Overall instruction to `setup_simulation`. Lists the names of the different files to call for each of the other folder categories, as well as the overall simulation parameters of name (a string), time (in hours as a float), and steps (number as an integer).

- **exchange_reactions:** Lists the exchange reactions that will be kept track of, giving both their ID and full name as seen in the model, as well as specifying the **initial_ub** and **initial_lb** and whether they are **limiting**. The **initial** parameters specify the absolute upper and lower bounds for the different exchange reactions, allowing them to be locked independently of fermentor conditions. If the **limiting** parameter is set to **true**, the amount of the corresponding metabolite in the medium will be used to limit the corresponding uptake reactions when it runs out (it does not set the max flux otherwise!), while if it is set to **false**, the amount of the metabolite will not be used to limit the corresponding uptake reaction. Note that the amount of the metabolite in the medium will still be tracked, and may become negative if the **limiting** parameter is set to **false**. This is by design, as it is useful to investigate hypotheticals, such as how much of the compound the model will use before something else potentially becomes limiting. Only the reaction ID column will be used to match with the model reactions. This file is automatically generated for a given model when the function `generate_setup_files` is called, after which it should be manually inspected and altered if needed.
- **media:** Contains the concentration (in $\mathrm{mmol\,L^{-1}}$) of different compounds in the medium, listed with reaction IDs and full names. Several of these files may be called in a single simulation.
- **models:** Subfolders containing the models in different formats.
  - **mat:** MATLAB files stored of the different model structures to allow for more rapid loading between simulations, as the COBRA Toolbox `readCbModel` function can be time-consuming. When run, `setup_simulation` looks in this folder for the model first, then looks for an SBML file if it cannot be found. They are then automatically generated and added to the **mat** folder when an SBML file containing one of the models is loaded for the first time.
  - **xml:** Models stored in the SBML format. Will only be searched if no appropriately-named MAT file is found first.

- **output:** Folder containing folders for the outputs of different simulations, bunched on a per-**input/configuration/** sub-file basis. files are added here both from `setup_simulation` and `plot_simulation`. The folder can be empty before a simulation is begun, as the program will automatically generate the necessary subfolders when required.
  - **<sim_name>:** The immediate subfolders in the **output/** folder will be automatically named after the corresponding file in **input/configuration/**.
    - **csv:** Comma-separated variable file (using semicolons as delimiters by default), containing some key parameters of interest for plotting.
    - **fig:** MATLAB FIG files for the plots generated by the `plot_simulation` function for the different simulations belonging to the corresponding file in **input/configuration/**.
    - **mat:** MATLAB mat files for the resulting MATLAB structs generated by the `run_simulation` function for the different simulations belonging to the corresponding file in **input/configuration/**. They are named by their configuration entry.
    - **png:** Images in the PNG format for the plots generated by the `plot_simulation` function for the different simulations belonging to the corresponding file in **input/configuration/**.

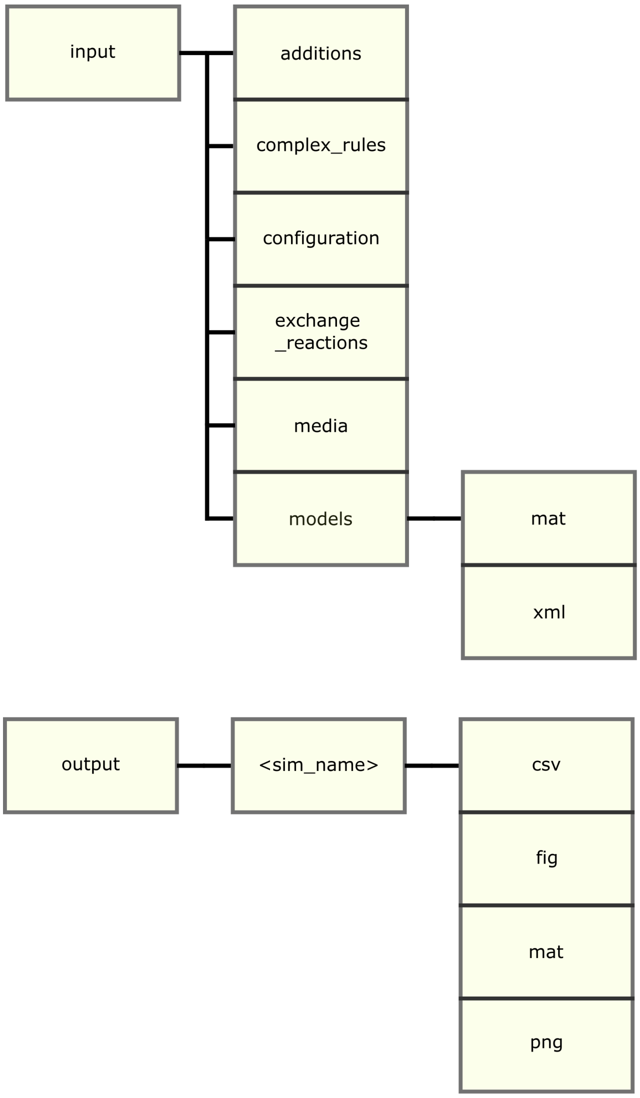


Figure 1: **Overview of file hierarchy.** An overview of the file hierarchy. Blocks located above one another are on the same level, blocks connected laterally by lines indicate a folder-subfolder relationship. Folder with variable names are named descriptively and marked with ”<” and ”>”.

### Program structure

The overall program structure is represented schematically in Fig. 2. The `setup_simulation` function is called with a string, specifying the name of a given csv configuration file (without the .csv extension). It loads the specified file from **input/configuration/**, reading in the various configuration parameters for each given simulation run, of which there can be several, one on each line. The parameters, for each simulation, are: the declared name of the simulation (**simulation_name**), the time in hours that the simulation should last for (**simulation_time_hours**), the number of simulation steps should happen in the given time frame (**simulation_time_step_number**), the amount of biomass that the fermentor simulation should be initiated with (**simulation_initial_biomass_gDW**), the name of the model file (**model_name**), the name of the biomass objective function that should be used (**biomass_function**), the name of the file specifying the exchange reactions (**exchange_reactions_file**), the name of the file specifying the media files (**media_files**), the name of the file specifying the additions program(**addition_program**), the name of the file specifying any custom complex rules (**complex_rules**), the amount of enzyme that can be redistributed per hour of simulation time (**enzyme_change_grams_per_hour**), and whether a summary plot should pop up after each simulation (**plot_live**).

The `setup_simulation` function builds a data structure containing all the pertinent information and passes this to the `run_simulation` function, which then performs the simulation as specified in the respective setup files. This is performed separately for each line in the file found in **input/configuration/**, so several different combinations of the files in the subfolders of **input/** can be used for several different simulations for one call to `setup_simulation`.

Each time it is run, `run_simulation` produces an output variable, which is a struct with all the input, as well as all the logged output information from the simulation run. That result is then saved as a MAT file, and is passed to `plot_simulation`, which produces a plot for the entire simulation duration, with time (in h) on the horizontal axis and growth (in $\mathrm{gCDW\,gCDW^{-1}\,h^{-1}}$), total biomass (in gCDW, and the total amount of all tracked external metabolite concentrations (in mmol) on the vertical axis. This is the summary plot that pops up directly if **plot_live** is set to 1.

Another plotting function, `plot_enzyme_distribution`, makes an area plot of enzyme pool utilization by different reactions against time. This is useful for visualizing how the total enzyme mass is distributed among the enzymes, and may be of assistance when troubleshooting strange model behavior.

#### The main loop

During the execution of `run_simulation`, after initializing some variables, the following steps are executed procedurally on repeat:

1. Update environment based on addition schedule
2. Update environment based on FBA sol
3. Determine if enzyme constraint if applicable
4. Limit uptake rates based on environment concentrations if applicable
5. Apply complex rules
6. Solve FBA
   (a) Solve initial FBA problem
   (b) Apply limits in enzyme composition if applicable and re-solve
   (c) If infeasible, perform bi-level optimization where growth rate is optimized subject to optimal values for demand sinks
7. Performing growth and perpetuating environment
   (a) Perform growth
   (b) Perpetuate environment and additions
   (c) Log data not already logged

A number of steps are executed as specified in the **input/configuration** file, before the result structure is returned to the `setup_simulation` script.

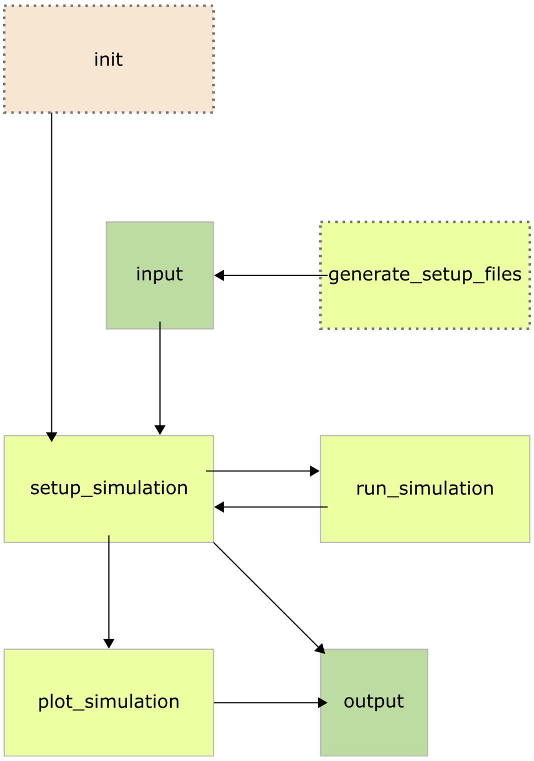


Figure 2: **Schematic of program flow.** A schematic representation of program flow. The function `generate_setup_files` can optionally be called to ease the preparation of the input data, and a suggested `init` script is included, but not necessary, to call the main program: `setup_simulation`. This main program then calls `run_simulation`, saves the return output in the **output/** folder, and sends it on to be plotted in `plot_simulation`. Here, the plotted figures are also saved in the **output/** folder.

#### Modeling the fermentor

The environment is handled as a perfectly mixed reservoir with no geometry and arbitrary capacity, which contains a given volume of total solution, and a given amount (mmol) of the different compounds in it. As components are added to, or removed from, the fermentation vessel, the total volume and the amounts of different compounds and components are updated accordingly. Concentrations are calculated when needed, by dividing the amount of a given compound by the total volume of fluid in the fermentor.

As it stands, the current implementation of the simulation environment is simple but serviceable Expansions by users are possible and encouraged.
