## Supplemental File 3 for "A study of a diauxic growth experiment using an expanded dynamic flux balance framework": generate_setup_files.m

```
function [] = generate_setup_files(model,outputName)
%generate_setup_files Helper script for fermentation simulator
%   This script accepts a COBRA model and a given output name, and
%   generates files required to run the simulation.


%% Generating columns
excInds = find(findExcRxns(model));
exchange_reaction_id = model.rxns(excInds);
exchange_reaction_name = model.rxnNames(excInds);
limiting = repmat("TRUE",length(excInds),1);
initial_ub = model.ub(excInds);
initial_lb = model.lb(excInds);
kineticZeroArray = zeros(size(initial_lb));
concentrations = zeros(length(excInds),1);

%% Generating exchange reactions file
varNames = ["exchange_reaction_id","exchange_reaction_name","limiting","initial_lb","initial_ub","vmax","km"];
outputTable = table(exchange_reaction_id,exchange_reaction_name,limiting,initial_lb,initial_ub,kineticZeroArray,kineticZeroArray,'VariableNames',varNames);
fileName = "input/exchange_reactions/"+outputName+".csv";
writetable(outputTable,fileName,'Delimiter',';');

%% Generating medium file
varNames = ["exchange_reaction_id","exchange_reaction_name","concentration_mmol_per_liter"];
outputTable = table(exchange_reaction_id,exchange_reaction_name,concentrations,'VariableNames',varNames);
fileName = "input/media/"+outputName+".csv";
writetable(outputTable,fileName,'Delimiter',';');

end
```
