## Supplemental File 3 for "A study of a diauxic growth experiment using an expanded dynamic flux balance framework": init.m

```
%% Initializing
% 
% AUTHOR:
%	- Emil Karlsen, March 2022
%
% NOTE:
%	This code is supplementary to the paper: A study of a diauxic growth
%	experiment using an expanded dynamic flux balance framework, Karlsen et
%	al. 2022
% 
%%%%%%%%%%%%%%%%%%%%%%%%%%%%%%%%%%%%%%%%%%%%%%%%%%%%%%%%%%%%%%%%
% % For optimizer parameters from paper, run the 2 lines below %
% changeCobraSolver('glpk')                                    %
% changeCobraSolverParams('LP', 'feasTol',1e-9)                %
%%%%%%%%%%%%%%%%%%%%%%%%%%%%%%%%%%%%%%%%%%%%%%%%%%%%%%%%%%%%%%%%

%% Reading settings

setup_simulation("CL_search");
setup_simulation("CL_ecc");
setup_simulation("CL_dFBA_compare");
```
