## Supplemental File 3 for "A study of a diauxic growth experiment using an expanded dynamic flux balance framework": plot_enzyme_distribution.m

```
function [] = plot_enzyme_distribution(fermStruct)


%% Plotting concentrations, growth, biomass, and volume

% Making some convenient variables
titleString = fermStruct.afterSimulation.settings.simulation_parameters.simulation_name+" enzyme distribution";

% Making figure
h = figure('Position',[100 100 1000 800]);
ax = axes;
nonZeroInds = sum(fermStruct.afterSimulation.results.enzyme_mass_distribution,2)~=0;
area(ax,fermStruct.afterSimulation.results.enzyme_mass_distribution(nonZeroInds,:)');

configFile = fermStruct.beforeSimulation.settings.configFileName;
[ status, msg ] = mkdir('output',char(configFile));
[ status, msg ] = mkdir(char("output/"+configFile),'png');
[ status, msg ] = mkdir(char("output/"+configFile),'fig');
savePath = char("output/"+configFile+"/fileType/");

figSaveString = replace(savePath,"fileType","png")+replace(titleString," ","_")+"_enzymeDist"+".png";
saveas(h,figSaveString);
figSaveString = replace(savePath,"fileType","fig")+replace(titleString," ","_")+"_enzymeDist"+".fig";
saveas(h,figSaveString);


end
```
