## Supplemental File 3 for "A study of a diauxic growth experiment using an expanded dynamic flux balance framework": plot_simulation.m

```
function [] = plot_simulation(fermStruct,yLims)
%PLOT_SIMULATION Used by setup_simulation to plot output data
%
% INPUT:
%    fermentation:	Matlab struct with configuration details
% 
% OPTIONAL INPUTS:
%    None
% 
% AUTHOR:
%	- Emil Karlsen, March 2022
%
% NOTE:
%	This code is supplementary to the paper A study of a diauxic growth
%	experiment using an expanded dynamic flux balance framework, Karlsen et
%	al. 2022
% 

%% Plotting concentrations, growth, biomass, and volume

% Setting fermentation variable for less verbose code
fermentation = fermStruct.afterSimulation;

% Finding interesting environmental concentrations
plotIndexes = find(sum(abs(fermentation.results.environment),2));
fprintf("%i environmental concentrations non-zero at any point during simulation\n",length(plotIndexes));
plotIndexNames = string(fermentation.settings.exchange_reactions{plotIndexes,1});
waterIndex = plotIndexNames=="EX_h2o_e";
plotIndexes(waterIndex) = [];
plotIndexNames(waterIndex) = [];

% Making some convenient variables
envDat = fermentation.results.environment(plotIndexes,:);
maxTime = fermentation.settings.simulation_parameters.simulation_time_hours;
multiplier = max(envDat,[],'all')/max(fermentation.results.totalBiomass);
if multiplier > max(envDat,[],'all')
    multiplier = 0.5*max(envDat,[],'all');
end
xAx = fermentation.results.time;
legendText = ["Biomass x"+string(multiplier); "Objective x"+string(multiplier); "Volume x"+string(multiplier); plotIndexNames];
titleString = fermentation.settings.simulation_parameters.simulation_name;

% Making irregular pattern of color, line style, and marker style
col = hsv(length(plotIndexes));
lineStyles = ["-","--",":","-."];
markers = ["o","+","*","x","^","v"];
styleIt = 1;
markerIt = 1;

% Making figure
plot_live = 'off';
if fermStruct.beforeSimulation.settings.simulation_parameters.plot_live == 1
    plot_live = 'on';
end
figure('Position',[100 100 1000 800],'visible',plot_live);
hold on;
plot(xAx,fermentation.results.totalBiomass*multiplier,'color','k','LineStyle','-','Marker',"x");
plot(xAx,fermentation.results.objective*multiplier,'color','k','LineStyle','--','Marker',"^");
plot(xAx,fermentation.results.volume*multiplier,'color','k','LineStyle','--','Marker',"o");
for i=1:length(plotIndexes)
    plot(xAx,envDat(i,:),'color',col(i,:),'LineStyle',lineStyles(styleIt),'Marker',markers(markerIt))
    styleIt = styleIt+1;
    markerIt = markerIt+1;
    if styleIt>length(lineStyles)
        styleIt = 1;
    end
    if markerIt>length(markers)
        markerIt = 1;
    end
end

% Making legend
legend(legendText,'Interpreter','none')
title(replace(titleString,"_"," "))

configFileName = fermStruct.beforeSimulation.settings.configFileName;
[ status, msg ] = mkdir('output',char(configFileName));
[ status, msg ] = mkdir(char("output/"+configFileName),'png');
[ status, msg ] = mkdir(char("output/"+configFileName),'fig');
savePath = char("output/"+configFileName+"/fileType/");

figHandle = gcf;
figSaveString = replace(savePath,"fileType","png")+replace(titleString," ","_")+"_metabolites"+".png";
saveas(figHandle,figSaveString);
figSaveString = replace(savePath,"fileType","fig")+replace(titleString," ","_")+"_metabolites"+".fig";
saveas(figHandle,figSaveString);

%% Plotting enzyme levels without legend to find stable compositions

if fermStruct.beforeSimulation.settings.enzyme_change_grams_per_hour
    figure('visible',plot_live)
    plot(fermStruct.afterSimulation.results.enzyme_mass_distribution');
    hold on
    plot(fermStruct.afterSimulation.results.massDistanceFromOptimal,'--');
    figHandle = gcf;
    figSaveString = replace(savePath,"fileType","png")+replace(titleString," ","_")+"_enzymes"+".png";
    saveas(figHandle,figSaveString);
    figSaveString = replace(savePath,"fileType","fig")+replace(titleString," ","_")+"_enzymes"+".fig";
    saveas(figHandle,figSaveString);
    xlabel("time")
    ylabel("enzyme levels (-), dist from opt (--)")
end

end
```
