## Supplemental File 3 for "A study of a diauxic growth experiment using an expanded dynamic flux balance framework": run_simulation.m

```
function fermentation = run_simulation(fermentation)
%RUN_SIMULATION Runs a simulation of growth in medium
%
% INPUT:
%    fermentation:	Matlab struct with configuration details
% 
% OPTIONAL INPUTS:
%    None
% 
% AUTHOR:
%	- Emil Karlsen, March 2022
%
% NOTE:
%	This code is supplementary to the paper: A study of a diauxic growth
%	experiment using an expanded dynamic flux balance framework, Karlsen et
%	al. 2022
% 

%% Loading simulation parameters

simulationIndex = 1;
simulationName = string(fermentation.settings.simulation_parameters.simulation_name);
simulation_time_hours = fermentation.settings.simulation_parameters.simulation_time_hours;
simulation_time_step_number = fermentation.settings.simulation_parameters.simulation_time_step_number;
cellBiomass = fermentation.settings.simulation_parameters.initial_biomass_gDW;
biomass_index = find(string(fermentation.settings.model.rxns)==fermentation.settings.simulation_parameters.biomass_function);

%% Initializing variables

% Volume
totalVolume = 0;
% Model
model = fermentation.settings.model;
sol = optimizeCbModel(model);
% AdditionCheck
checkAdditions = true;
% Exchange indexes
exchangeIndices = zeros(size(fermentation.settings.exchange_reactions.exchange_reaction_id));
for i=1:length(exchangeIndices)
    exchangeIndices(i) = find(model.rxns == string(fermentation.settings.exchange_reactions.exchange_reaction_id(i)));
end
% Enzyme tracking assuming sMOMENT
allowed_enz_change_on_step = nan;
fermentation.results.enzyme_mass_distribution = zeros(length(model.rxns)-1,simulation_time_step_number);

% Fetching enzyme mass per flux from model
% NB: This will only work for sMOMENT models, and has only been tested with
% the iJO1366* sMOMENT model
enzyme_indices = [true(length(model.rxns)-1,1);false]; %works with sMOMENT
enzyme_mass_per_flux = -model.S(end,enzyme_indices);

%% Some constants

time_hours_per_step = simulation_time_hours/simulation_time_step_number;
stepPrint = num2str(numel(num2str(simulation_time_step_number)));

%% Initializing results

fermentation.results.additions = zeros(height(fermentation.settings.additions),simulation_time_step_number);
fermentation.results.fluxes = zeros(length(model.rxns),simulation_time_step_number);
fermentation.results.optimalFluxes = zeros(length(model.rxns),simulation_time_step_number);
fermentation.results.enzymeFluxCapacities = zeros(length(model.rxns)-1,simulation_time_step_number);
fermentation.results.biomass = zeros(1,simulation_time_step_number);
fermentation.results.uptake_limits = zeros(length(exchangeIndices),simulation_time_step_number);
fermentation.results.volume = zeros(1,simulation_time_step_number);
fermentation.results.objective = zeros(1,simulation_time_step_number);
fermentation.results.time = zeros(1,simulation_time_step_number);
fermentation.results.ubs = zeros(length(model.rxns),simulation_time_step_number);
fermentation.results.lbs = zeros(length(model.rxns),simulation_time_step_number);
fermentation.results.cs = zeros(length(model.c),simulation_time_step_number);
fermentation.results.massDistanceFromOptimal = zeros(simulation_time_step_number,1);

%% Main loop:

for step_t=1:simulation_time_step_number
    time_at_step = step_t/simulation_time_step_number*simulation_time_hours;
    
    %% Updating environment based on addition schedule
    
    % Adding additions
    if checkAdditions
        allAdded = true;
        for i=1:length(fermentation.settings.additions.add_time_start_hours)
            if time_at_step >= fermentation.settings.additions.add_time_start_hours(i)
                % Add from media list; enable adding met directly?
                allAdded = false;
                added = fermentation.results.additions(i,step_t);
                toAdd = fermentation.settings.additions.add_amount_liters(i);
                if added < toAdd
                    difference = toAdd-added;
                    add_time_duration_hours = fermentation.settings.additions.add_time_duration_hours(i);
                    proportion_to_add = time_hours_per_step/add_time_duration_hours;
                    amount_to_add = toAdd*proportion_to_add;
                    if amount_to_add > difference
                        amount_to_add = difference;
                    end
                    try
                        if fermentation.settings.additions.medium_id{i} == "outflow"
                            % Handling of outflow "addition"
                            volumeProportionRemoved = amount_to_add/totalVolume;
                            volumeProportionRemaining = 1-volumeProportionRemoved;
                            if volumeProportionRemaining > 1 || volumeProportionRemaining < 0
                                error("%.2f\% of volume attempted removed in one time step!\n",volumeProportionRemaining*100);
                            end
                            for met=1:length(fermentation.results.environment(:,step_t))
                                fermentation.results.environment(met,step_t) = fermentation.results.environment(met,step_t)*volumeProportionRemaining;
                            end
                            cellBiomass = cellBiomass*volumeProportionRemaining;
                            totalVolume = totalVolume*volumeProportionRemaining;
                        else
                            % Handling of regular additions (media)
                            medTab = fermentation.settings.media.(string(fermentation.settings.additions.medium_id(i)));
                            for j=1:height(medTab)
                                environment_index = find(string(fermentation.settings.exchange_reactions.exchange_reaction_id)==string(medTab.exchange_reaction_id(j)));
                                metToAdd = amount_to_add*medTab.concentration_mmol_per_liter(j);
                                fermentation.results.environment(environment_index,step_t) = fermentation.results.environment(environment_index,step_t)+metToAdd;
                            end
                            % Increasing volume correspondingly
                            totalVolume = totalVolume+amount_to_add;
                            fermentation.results.additions(i,step_t) = fermentation.results.additions(i,step_t)+amount_to_add;
                        end
                    catch
                        warning("Error attempting addtion [%s]\n",string(fermentation.settings.additions{i,1}))
                    end
                end
            end
        end
        if allAdded
            checkAdditions = false;
        end
    end
    
    %% Updating environment based on FBA sol
    
    % Counting and listing negative concentrations before FBA update
    negativeEnvironmentConcentrations = length(find(fermentation.results.environment(:,step_t)<-0.01));
    if negativeEnvironmentConcentrations
        fprintf("%i reactions at negative concentration before fba subtraction\n",negativeEnvironmentConcentrations);
    end
    
    % Modifying environment based on FBA solution if not initial step
    if step_t>1
        lastGR = fermentation.results.fluxes(biomass_index,step_t-1);
        lastBM = fermentation.results.totalBiomass(step_t-1);
        meanBM = lastBM/(time_hours_per_step*lastGR)*(exp(lastGR*time_hours_per_step)-1);
        if isnan(meanBM)
            meanBM = lastBM;
        end
        fermentation.results.environment(:,step_t) = fermentation.results.environment(:,step_t)+sol.x(exchangeIndices)*time_hours_per_step*meanBM;
    end
    
    % Counting and listing negative concentrations after FBA update
    negativeEnvironmentConcentrations = length(find(fermentation.results.environment(:,step_t)<-0.01));
    if negativeEnvironmentConcentrations
        fprintf("%i reactions at negative concentration after fba subtraction\n",negativeEnvironmentConcentrations);
    end
    
    
    %% Determine enzyme constraint if applicable
    
    if istable(fermentation.settings.enzyme_change_grams_per_hour)
        for row=1:height(fermentation.settings.enzyme_change_grams_per_hour)
            var = string(fermentation.settings.enzyme_change_grams_per_hour.variable{row});
            varInd = find(string(model.rxns(:))==var);
            assert(length(varInd) == 1);
            con = fermentation.settings.enzyme_change_grams_per_hour.condition_for_variable(row);
            varVal = fermentation.settings.enzyme_change_grams_per_hour.value_of_variable(row);
            if con == "geq"
                if fermentation.results.fluxes(varInd) >= varVal
                    allowed_enz_change_on_step = time_hours_per_step*fermentation.settings.enzyme_change_grams_per_hour.enzyme_change_grams_per_gDW_per_hour(row);
                end
            end
        end
    elseif fermentation.settings.enzyme_change_grams_per_hour
        allowed_enz_change_on_step = time_hours_per_step*fermentation.settings.enzyme_change_grams_per_hour;
    end
   
    
    %% Limit uptake rates based on environment concentrations
    for met=1:length(exchangeIndices)
        % Simple "all available all the time" law, except where km is given
%         availableMet = fermentation.results.environment(met,step_t)/(cellBiomass*exp(time_hours_per_step*sol.f)*time_hours_per_step);
        availableMet = fermentation.results.environment(met,step_t)/(cellBiomass*time_hours_per_step);
        if fermentation.settings.exchange_reactions.vmax(met) ~= 0 && fermentation.settings.exchange_reactions.km(met) ~= 0
            vmax = fermentation.settings.exchange_reactions.vmax(met);
            km = fermentation.settings.exchange_reactions.km(met);
            subConc = fermentation.results.environment(met,step_t);
            mmLim = vmax*subConc/(km+subConc);
            if mmLim<availableMet
                availableMet = mmLim;
            end
        end
        if availableMet < 0
            availableMet = 0;
        end
        % Reporting spent resources
        if fermentation.settings.exchange_reactions.limiting{met}
            if model.lb(exchangeIndices(met))<0 && availableMet==0
                fprintf("Ran out of %s!\n",string(model.rxns(exchangeIndices(met))));
            end
            model.lb(exchangeIndices(met)) = -availableMet;
        end
        if model.lb(exchangeIndices(met)) < fermentation.settings.exchange_reactions.initial_lb(met)
            model.lb(exchangeIndices(met)) = fermentation.settings.exchange_reactions.initial_lb(met);
        end
        if model.ub(exchangeIndices(met)) > fermentation.settings.exchange_reactions.initial_ub(met)
            model.ub(exchangeIndices(met)) = fermentation.settings.exchange_reactions.initial_ub(met);
        end
    end
    fermentation.results.uptake_limits(:,step_t) = model.lb(exchangeIndices);
    
    
    %% Apply complex rules
    
    if istable(fermentation.settings.complex_rules)
        for i=1:height(fermentation.settings.complex_rules)
            constraint = "";
            rule = fermentation.settings.complex_rules.rule(i);
            k1 = fermentation.settings.complex_rules.k1(i);
            k2 = fermentation.settings.complex_rules.k2(i);
            scalingFactor = 1;
            if contains(fermentation.settings.complex_rules.constraint(i),"lb.")
                constraint = replace(fermentation.settings.complex_rules.constraint(i),"lb.","");
            elseif contains(fermentation.settings.complex_rules.constraint(i),"ub.")
                constraint = replace(fermentation.settings.complex_rules.constraint(i),"ub.","");
            end
            metExcIndex = find(string(fermentation.settings.exchange_reactions.exchange_reaction_id) == constraint);
            index = find(string(model.rxns) == constraint);
            concentration = fermentation.results.environment(metExcIndex, step_t)/totalVolume;
            if rule == "scale_by_delta_linear"
                scalingFactor = (k1-concentration)/k1;
            elseif rule == "scale_by_delta_exponetial"
                scalingFactor = (exp(k1)-exp(concentration))/exp(k1);
            elseif rule == "scale_by_ln"
                if concentration >= k1
                    alpha = 1/log(k2/k1);
                    beta = 1/k2;
                    scalingFactor = -alpha*log(concentration*beta);
                elseif concentration >= k2
                    scalingFactor = 0;
                end
            end
            if contains(fermentation.settings.complex_rules.constraint(i),"lb.")
                model.lb(index) = fermentation.settings.exchange_reactions.initial_lb(metExcIndex)*scalingFactor;
            elseif contains(fermentation.settings.complex_rules.constraint(i),"ub.")
                model.ub(index) = fermentation.settings.exchange_reactions.initial_ub(metExcIndex)*scalingFactor;
            else
                warning("No bound found\n");
            end
        end
    end
    
    
    %% Solve FBA based on environment and log
    tempModel = model;
    sol = optimizeCbModel(tempModel);
    infeasibleCounter = 0;
    if isempty(sol.x)
        infeasibleCounter = infeasibleCounter+1;
        % If model infeasible, bi-level optimization
        demand_sinks = tempModel.lb;
        demand_sinks(tempModel.lb<=0) = 0;
        tempModel.c = demand_sinks;
        tempModel.lb(demand_sinks>0) = 0;
        sol = optimizeCbModel(tempModel);
        % Bi-level optimization
        demand_indexes = find(demand_sinks>0);
        tempModel = model;
        numTol = 10e-4;
        tempModel.lb(demand_indexes) = min(sol.x(demand_indexes)-numTol,0);
        sol = optimizeCbModel(tempModel);
        %
        fermentation.results.fluxes(:,step_t) = sol.x;
        assert(length(sol.x)>1);
    else
        fermentation.results.optimalFluxes(:,step_t) = sol.x;
    end
    
    % Adjust FBA for enzyme change constraint if applicable
    if ~isnan(allowed_enz_change_on_step)
        if step_t > 1
            enzyme_dist_targ = enzyme_mass_per_flux'.*sol.x(enzyme_indices);
            enzyme_dist_curr = fermentation.results.enzyme_mass_distribution(:,step_t-1);
            mass_reallocation_vector = enzyme_dist_targ-enzyme_dist_curr;
            mass_addition_vector = mass_reallocation_vector;
            mass_addition_vector(mass_addition_vector<0) = 0;
            mass_addition_sum = sum(mass_addition_vector);
            mass_reallocation_proportion_possible = min(1,allowed_enz_change_on_step/mass_addition_sum);
            enzyme_dist_new = enzyme_dist_targ*mass_reallocation_proportion_possible+enzyme_dist_curr*(1-mass_reallocation_proportion_possible);
            fermentation.results.enzyme_mass_distribution(:,step_t) = enzyme_dist_new;
            for enz=1:length(enzyme_dist_new)
                if enzyme_mass_per_flux(enz) > 0
                    tempModel.ub(enz) = enzyme_dist_new(enz)/enzyme_mass_per_flux(enz);
                    if ~(tempModel.ub(enz) >= 0)
                        tempModel.ub(enz) = 0;
                    end
                    assert(~isnan(tempModel.ub(enz)))
                end
            end
            fermentation.results.massDistanceFromOptimal(step_t) = sum(abs(mass_reallocation_vector));
            sol = optimizeCbModel(tempModel);
        else
            assert(step_t==1)
            fermentation.results.enzyme_mass_distribution(:,step_t) = enzyme_mass_per_flux'.*sol.x(enzyme_indices);
        end
    end
    
    if isempty(sol.x)
        infeasibleCounter = infeasibleCounter+1;
        if infeasibleCounter == 1
            warning("infeasible after enzyme limitations")
        elseif infeasibleCounter > 1
            error("infeasibleCounter triggered twice!");
        end
        % If model infeasible, remove objective and optimize for demand
        % sinks (usually just ATP maintenance) instead
        demand_sinks = tempModel.lb;
        demand_sinks(tempModel.lb<=0) = 0;
        tempModel.c = demand_sinks;
        tempModel.lb(demand_sinks>0) = 0;
        sol = optimizeCbModel(tempModel);
        assert(length(sol.x)>1);
    end
    
    % Save FBA results
    fermentation.results.fluxes(:,step_t) = sol.x;
    
    
    %% Performing growth, perpetuating, and logging extras
    
    % Perform growth and log it
    cellBiomass = cellBiomass*exp(time_hours_per_step*sol.x(biomass_index));
    fermentation.results.totalBiomass(step_t) = cellBiomass;
    fermentation.results.biomassPerVol(step_t) = cellBiomass/totalVolume;
    
    fermentation.results.objective(step_t) = sol.f;
    updateString = "Step: %"+stepPrint+"i, Biomass: %.4f, Growth: %.3f, Objective: %.3f, Volume (L): %.3f, allowed_enz_change_on_step: %.6f, massDistFromOpt: %.3f\n";
    fprintf(updateString,step_t,cellBiomass,sol.x(biomass_index),sol.f,totalVolume,allowed_enz_change_on_step,fermentation.results.massDistanceFromOptimal(step_t));
    
    % Perpetuate environment and additions
    if step_t < simulation_time_step_number
        fermentation.results.environment(:,step_t+1) = fermentation.results.environment(:,step_t);
        fermentation.results.additions(:,step_t+1) = fermentation.results.additions(:,step_t);
    end
    
    % Log extras
    fermentation.results.volume(step_t) = totalVolume;
    fermentation.results.time(step_t) = time_at_step;
    fermentation.results.ubs(:,step_t) = tempModel.ub;
    fermentation.results.lbs(:,step_t) = tempModel.lb;
    fermentation.results.cs(:,step_t) = tempModel.c;
    
    
end

end
```
