## Supplemental File 3 for "A study of a diauxic growth experiment using an expanded dynamic flux balance framework": setup_simulation.m

```
function fermStruct = setup_simulation(configFileName)
%SETUP_SIMULATION Reads configuration files before performing decFBAecc
%
% INPUT:
%    configFileName:	name of the configuration file to be used
% 
% OPTIONAL INPUTS:
%    None
% 
% AUTHOR:
%	- Emil Karlsen, March 2022
%
% NOTE:
%	This code is supplementary to the paper A study of a diauxic growth
%	experiment using an expanded dynamic flux balance framework, Karlsen et
%	al. 2022
% 

%% Printing configuration parameters

fprintf("Loading configuration table from %s\n",configFileName);
configTable = readtable(char("input/configuration/"+configFileName+".csv"),'Delimiter',';');
fprintf("Running %i simulations, with a total of %i steps\n",height(configTable),sum(configTable.simulation_time_step_number))
configTable
pause(2)

time = now;
timeString = datetime(time,'ConvertFrom','datenum','Format','yyyyMMdd_HH_mm_ss');

for simulationIterator=1:height(configTable)

%% Initial setup for simulation
    
fprintf("\n\n--------------------------------------------------------------------\n");
fprintf("Preparing simulation %i/%i\n",simulationIterator,height(configTable));

fprintf("Simulation parameters:\n");
fprintf("Name:                 %s\n",string(configTable.simulation_name(simulationIterator)));
fprintf("Time (h):             %s\n",string(configTable.simulation_time_hours(simulationIterator)));
fprintf("Steps:                %s\n",string(configTable.simulation_time_step_number(simulationIterator)));
fprintf("Initial BM:           %s\n",string(configTable.simulation_initial_biomass_gDW(simulationIterator)));
fprintf("Model:                %s\n",string(configTable.model_name(simulationIterator)));
fprintf("Exchange config file: %s\n",string(configTable.exchange_reactions_file(simulationIterator)));
fprintf("Media files:          %s\n",string(configTable.media_files(simulationIterator)));
fprintf("Addition program:     %s\n",string(configTable.addition_program(simulationIterator)));
fprintf("Complex rules:        %s\n",string(configTable.complex_rules(simulationIterator)));
fprintf("Enzyme change rate:   %s g/h\n",string(configTable.enzyme_change_grams_per_hour(simulationIterator)));
fprintf("Print live:           %s\n",string(configTable.plot_live(simulationIterator)));
fprintf("\n");

pause(1)

media = string(split(configTable.media_files(simulationIterator),','));
addition_program = string(configTable.addition_program(simulationIterator));
model_name = string(configTable.model_name(simulationIterator));
complex_rules = string(configTable.complex_rules(simulationIterator));
exchange_reactions = string(configTable.exchange_reactions_file(simulationIterator));
enzyme_change_grams_per_hour = string(configTable.enzyme_change_grams_per_hour(simulationIterator));
plot_live = configTable.plot_live(simulationIterator);

simulation_parameters = struct();
simulation_parameters.simulation_name = string(configTable.simulation_name(simulationIterator));
simulation_parameters.initial_biomass_gDW = configTable.simulation_initial_biomass_gDW(simulationIterator);
simulation_parameters.simulation_time_hours = configTable.simulation_time_hours(simulationIterator);
simulation_parameters.simulation_time_step_number = configTable.simulation_time_step_number(simulationIterator);
simulation_parameters.biomass_function = configTable.biomass_function(simulationIterator);
simulation_parameters.plot_live = plot_live;

fprintf("Reading simulation settings\n\n")
fermentation = struct();

%% Fetching configuration files

% Specifying source config file
fermentation.settings.configFileName = configFileName;

% Fetching media:
for i=1:length(media)
    locationChar = char("input/media/"+media(i)+".csv");
    fermentation.settings.media.(media(i)) = readtable(locationChar, 'Delimiter', ';');
end

% Fetching addition program m9_batch_fermentation:
locationChar = char("input/additions/"+addition_program+".csv");
fermentation.settings.additions = readtable(locationChar, 'Delimiter', ';');

% Fetching complex rules:
if ~(complex_rules == "none")
    locationChar = char("input/complex_rules/"+complex_rules+".csv");
    fermentation.settings.complex_rules = readtable(locationChar, 'Delimiter', ';');
else
    fermentation.settings.complex_rules = false;
end

% Fetching simulation parameters:
fermentation.settings.simulation_parameters = simulation_parameters;

% Fetching exchange reactions
locationChar = char("input/exchange_reactions/"+exchange_reactions+".csv");
fermentation.settings.exchange_reactions = readtable(locationChar, 'Delimiter', ';');

% Fetching enzyme change limit
if str2double(enzyme_change_grams_per_hour) == -1
    fermentation.settings.enzyme_change_grams_per_hour = false;
else
    if ~isempty(str2double(enzyme_change_grams_per_hour))
        fermentation.settings.enzyme_change_grams_per_hour = str2double(enzyme_change_grams_per_hour);
    else
        locationChar = char("input/enzyme_change_grams_per_hour/"+enzyme_change_grams_per_hour+".csv");
        fermentation.settings.enzyme_change_grams_per_hour = readtable(locationChar, 'Delimiter', ';');
    end
end

clear locationChar


%% Sanity checks

% Checking that all listed media are present
fn = fieldnames(fermentation.settings.media);
additions = fermentation.settings.additions{:,1};
warned = false;
for i=1:length(additions)
    if ~sum(contains(fn,additions{i}))~=0 && additions{i}~="outflow"
        warning("Missing medium: [%s]!\n",additions{i})
        warned = true;
    end
end

% Listing available media in the case that not all listed are available
if warned
    fprintf("Available media are:\n")
    for i=1:length(fn)
        fprintf("%s\n",string(fn(i)));
    end
    fprintf("\n");
end

% Cleanup
clear fn additions i warned


%% Loading model

disp("Loading model")
loadPath = "input/models/";

try
    model = load(char(loadPath+"mat/"+model_name));
    model = model.model;
catch
    model = readCbModel(char([loadPath+"xml/"+model_name+".xml"]));
    save(char(loadPath+"mat/"+model_name),'model');
end

fermentation.settings.model = model;

% Opening all exchange reactions
fermentation.settings.exchangeIndices = find(contains(string(fermentation.settings.model.rxns),string(fermentation.settings.exchange_reactions.exchange_reaction_id)));
fermentation.settings.model.ub(fermentation.settings.exchangeIndices) = fermentation.settings.exchange_reactions.initial_ub;

% Cleanup
clear model_name structName loadPath


%% Setting up environment

disp("Setting up environment")

exchangeReactions = string(fermentation.settings.exchange_reactions.exchange_reaction_id);
for i=1:length(fermentation.settings.exchange_reactions.limiting)
    if string(fermentation.settings.exchange_reactions.limiting{i}) == "FALSE"
        fermentation.settings.exchange_reactions.limiting{i} = false;
    elseif string(fermentation.settings.exchange_reactions.limiting{i}) == "TRUE"
        fermentation.settings.exchange_reactions.limiting{i} = true;
    else
        error("Non-boolean statement in exchange_reactions limiting column")
    end
end
environment = cell(length(exchangeReactions),fermentation.settings.simulation_parameters.simulation_time_step_number);
fermentation.results.environment = zeros(size(environment));

% Cleanup
clear exchangeReactions environment


%% Running simulation

% Setting up save string and superstruct
fermStruct = struct();
fermStruct.beforeSimulation = fermentation;
saveName = replace(string(fermentation.settings.simulation_parameters.simulation_name)," ","_");
fprintf("Running simulation: %s\n\n",saveName)

% Running
fermentation = run_simulation(fermentation);

% Preparing to save
negInds = find(fermentation.results.environment(:,end)<0);
fermentation.results.negativeTerminalConcentrations = [fermentation.settings.exchange_reactions.exchange_reaction_id(negInds), string(fermentation.results.environment(negInds,end))];
fermStruct.afterSimulation = fermentation;

fprintf("%i environmental concentrations negative at end\n",length(negInds));

% Save and cleanup
[ status, msg ] = mkdir('output',char(configFileName));
[ status, msg ] = mkdir(char("output/"+configFileName),'mat');
savePath = char("output/"+configFileName+"/mat/"+saveName);
save(savePath,'fermStruct');
clear negInds

%% Saving data in CSV file

outputTable = table;
outputTable.Time = fermStruct.afterSimulation.results.time';
outputTable.TotalBiomass = fermStruct.afterSimulation.results.totalBiomass';
outputTable.BiomassPerVol = fermStruct.afterSimulation.results.biomassPerVol';
outputTable.Objective = fermStruct.afterSimulation.results.objective';
outputTable.Volume = fermStruct.afterSimulation.results.volume';
model = fermStruct.afterSimulation.settings.model;
for rxn=1:length(model.rxns)
    fn = string(model.rxns(rxn));
    dat = fermStruct.afterSimulation.results.fluxes(rxn,:)';
    outputTable.(fn) = dat;
end
for exc=1:size(fermStruct.afterSimulation.results.environment,1)
    index = fermStruct.afterSimulation.settings.exchangeIndices(exc);
    fn = "EnvironmentAmount_"+string(model.rxns(index));
    dat = fermStruct.afterSimulation.results.environment(exc,:)';
    outputTable.(fn) = dat;
end

[ status, msg ] = mkdir(char("output/"+configFileName),'csv');
savePath = char("output/"+configFileName+"/csv/"+saveName+".csv");
writetable(outputTable,savePath,'Delimiter',';');

%% Plotting results

plot_simulation(fermStruct,[0,50]);

end

end
```
